# AIM-CICs: automatic identification method for Cell-in-cell structures based on convolutional neural network

**DOI:** 10.1101/2021.02.26.432996

**Authors:** Meng Tang, Yan Su, Wei Zhao, Zubiao Niu, Banzhan Ruan, Qinqin Li, You Zheng, Chenxi Wang, Yong Zhou, Bo Zhang, Fuxiang Zhou, Hongyan Huang, Hanping Shi, Qiang Sun

## Abstract

Whereas biochemical markers are available for most types of cell death, current studies on non-autonomous cell death by entosis relays strictly on the identification of cell-in-cell structure (CICs), a unique morphological readout that can only be quantified manually at present. Moreover, the manual CICs quantification is generally over-simplified as CICs counts, which represents a major hurdle against profound mechanistic investigations. In this study, we take advantage of artificial intelligence (AI) technology to develop an automatic identification method for CICs (AIM-CICs), which performs comprehensive CICs analysis in an automated and efficient way. The AIM-CICs, developed on the algorithm of convolutional neural network (CNN), can not only differentiate between CICs and non-CICs (AUC > 0.99), but also accurately categorize CICs into five subclasses based on CICs stages and cell number involved (AUC > 0.97 for all subclasses). The application of AIM-CICs would systemically fuel researches on CICs-mediated cell death such as high-throughput screening.

## Introduction

Cell-in-cell structures (CICs) typically referred to the unusual eukaryotic cells involving the whole objects internalized partially or completely inside of others, which had been observed in diverse physiological and pathological samples [1, 2]. The presence of CICs was reported to be correlated with patient prognosis in a group of human tumors, such as breast cancer [3], head and neck squamous carcinoma [4, 5], and pancreatic ductal adenocarcinoma [6]. Functional studies implicated CICs in a number of biomedical processes, including embryonic development [7], mitotic surveillance [8], tumor evolution [9], and immune homeostasis [10] and the forth. As an evolutionarily conserved process, CICs formation was underlain by multiple mechanisms, such as entosis [11], cannibalism [12] and emperitosis [13]. Among which, entosis was one of the best studied processes that generally ended up with the death of the internalized cells in an acidified lysosome-dependent way [11, 14]. The formation of entotic CICs turned out to be a genetically controlled process, where cell internalization was driven cell-autonomously by polarized actomyosin resulted from the E-cadherin-mediated adherens junctions [15, 16], and coordinated by a mechanical ring interfacing in between them [17]. Additionally, an ever-expanding set of factors, acting through either actomyosin, or adherent junctions or mechanical ring, were identified as important regulators [18, 19, 20, 21].

Despite great progress made over the past decade, the studies on CICs formation were, however, based on the over-simplified readout of CICs counts that was performed manually, which is not only labor-intensive and time-consuming, but also sharply incompatible with the complex CICs formation per se. First, since CICs formation is a dynamic process preceding through sequential steps including cell-cell contact, penetration and closing [22], therefore, it generally gives rise to CICs at different stages displaying morphologies of partial or complete. Second, the CICs morphologies were further complicated by the involvement of multiple cells, which frequently resulted in structures of “cell-in-cell-in-cell” or even more. Third, due to personal experiences and preferences, the CICs judgment and inclusion-exclusion criteria for analysis varied from investigators to investigators, making it hard to compare across studies from different labs, or even studies from different investigators in one lab. In addition, manual quantification is rather inefficient in dealing with a large number of samples that may serve the screening purpose. Thus, the traditional CICs quantification reported results of less informative, hardly comparable and low-throughput, which calls for more efficient and informative ways for the quantification of CICs.

Recent years had witnessed the rapid development of image-based artificial intelligence (AI) technology in assisting biomedical practices. For example, by using a single convolutional neural networks (CNN) algorithm, Esteva *et al* demonstrated the classification of skin lesions in performance on par with all tested experts [23]. Lin *et al* developed a ResNeXt WSL model that achieved impressive performance (94.09% accuracy, 92.79% sensitivity, and 98.03% specificity) in making chromosome cluster type identification [24]. Actually, simply based on microscopic images, AI algorithms were quite competent in analyzing most, if not all, biological events such as the early onset of pluripotent stem cell differentiation [25], tumor cell malignancy [26], mitosis staging [27], and the like. The remarkable potentials in accuracy and efficiency make AI-based image analysis an ideal method for comprehensive and reliable CICs quantification.

In this study, based on RGB fluorescent microscopic images, we employed the deep CNN algorithms (Faster-RCNN and ResNet) to evaluate a large amount of cell candidates with defined subtypes and trained a multiclassfier for the recognition of subdivided CICs, which was named as AIM-CICs abbreviated from Automatic Identification Method of Cell-In-Cell structures. The AIM-CICs exhibited a high level of sensitivity and specificity, as evidenced by AUC values of > 0.97 for all tasks, in differentiating CICs from non-CICs, and identifying subtyped CICs from multiple cells. The development and application of AIM-CICs hold the promise of speeding up CICs-related studies, such as deciphering the molecular controls of CICs formation in a finer resolution, and enabling image-based systemic screening by high-content microscopy.

## Results

### The deep-learning framework of AIM-CICs

In this work, we conducted a framework of object detection and classification based on manual annotation in the training and validation set, and then performed inspections in the test set (Fig. 1). For an RGB-format image, the proposed system performs two consecutive steps. First, a Faster-RCNN [28] network with ResNet-50 [29] backbone was formulated to find the cell regions and extract the candidate patches. Second, each candidate, representing one cell or CICs, was classified by an ResNet-101 network based on the cellular morphology. Subsequently, those subdivided candidates of the predicted results were grouped into different folders, and marked out on the original locations of the corresponding images.

**Fig. 1.**
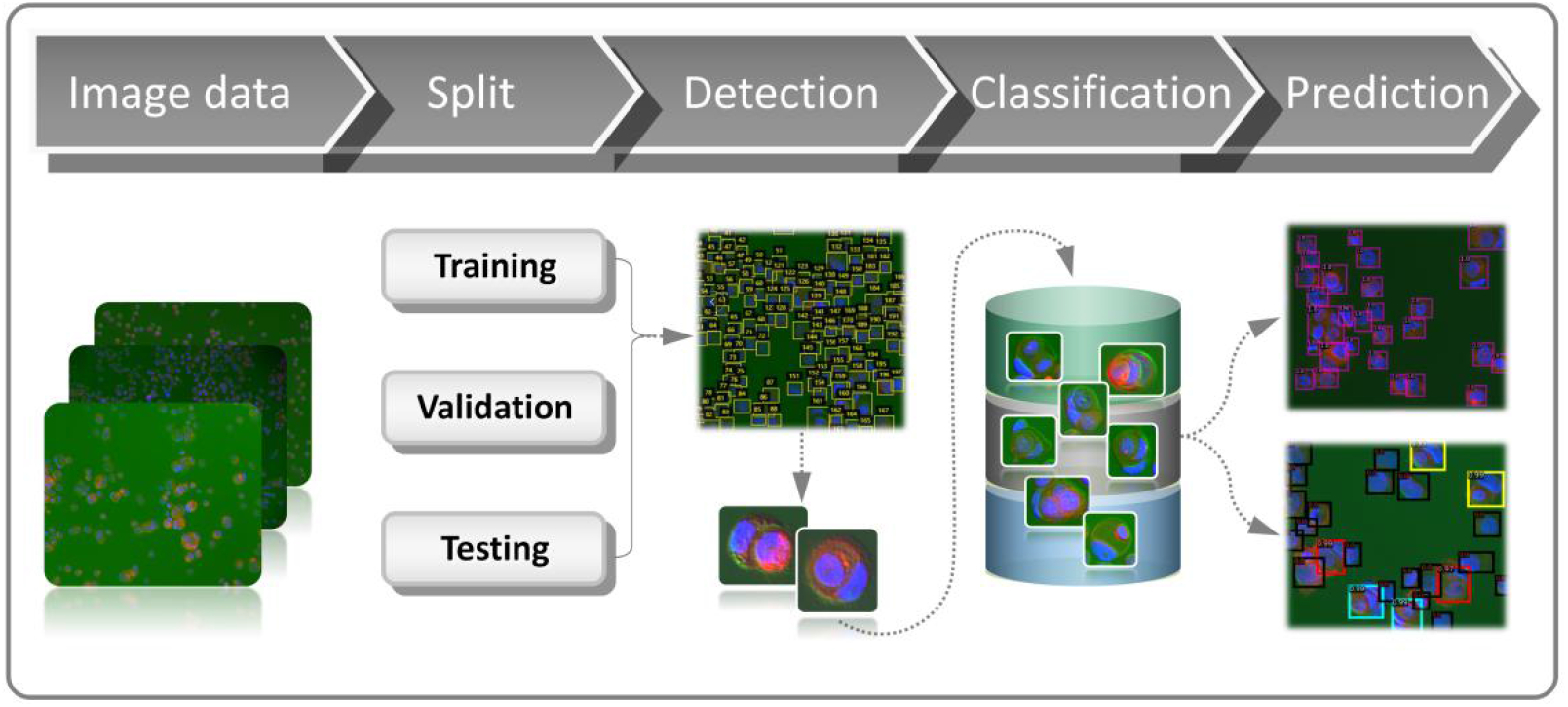
The workflow for the development of AIM-CICs.

### Cell region detection and extraction

Cell region detection is the initial task to investigate microscopic images. According to the basic cell components, we acquired the fluorescent microscopic images with red channel for membrane and blue channel for nucleus. Along with the bright field, the merged images could be further composited into RGB format with variant cell quantities and brightness values (Fig. S1a). The extraction of cell candidate aims to propose regions of interest (ROI) that potentially involved CIC structures. This step served to reduce the searching space and improve efficiency of subsequent steps in a high-content study. Initially, four pieces of MCF7 images and four pieces of MCF10A images, which included 2164 cells in total, were used as the training set for cell region detection. Through manually annotating these images using VGG Image Annotator (VIA, https://www.robots.ox.ac.uk/~vgg/software/via/) (Fig. 2a), cell region detection was further treated as a classic 1-class object detection task through the Faster-RCNN [28] network with ResNet-50 [29] backbone. Specifically, during training, we have performed random flip, random rotation and random scale for data augmentation, which greatly expanded the data’s diversity. Following the training process (Fig. 2b), we ensured the applicability of this step with average of 0.88 precision and 0.96 recall (IoU 0.1) by randomly testing on 10 pieces of MCF7 and MCF10A images, which covered 2398 cells (Fig. 2c).

**Fig. 2.**
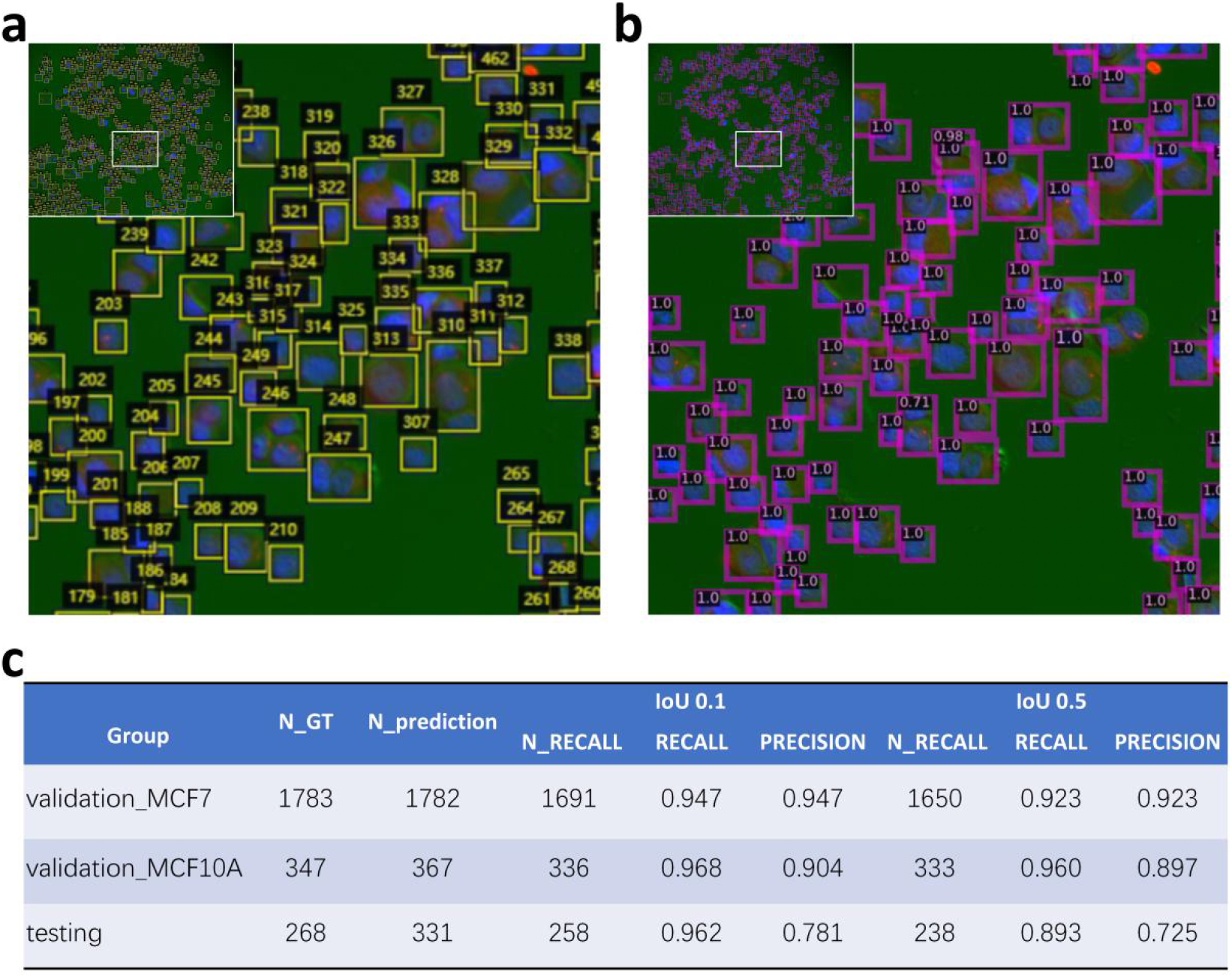
Cell regions detection and extraction. **a** Using VGG Image Annotator to annotate cell regions in the training set. Each box represents an individual cell region with numeric order on it. **b** Detection Model-based prediction of the cell region in the validation set. Each cell region was indicated by a box with a predicted confidence score. **c** Performance of cell region detection model. The MCF7 and MCF10A images in the validation set belong to the same batch of training set images, and test set is composed of MCF7 and MCF10A images from independent experiments. N_GT: number of ground truth cells. N_prediction: cell number of model’s prediction (confidence threshold set to 0.1). IoU: Intersection over Union.

It is believed that factors, such as cell morphology, sample density, as well as image brightness, do impact the accuracy of target detection and recognition. In the data collected this study, MCF10A samples generally displayed a larger cell size and much more complicated pattern of CICs as compared with MCF7 samples (Fig. S1a). Based on the precisely manual labeling, we could minimize the effect of target varieties among MCF7 and MCF10A samples (Fig. S1b), except for the over-exposed fluorescent images that should be excluded in the processing of the primary images. Eventually, we exported the patches of detected cell regions of the entire RGB-format images for the following analysis.

### Definition of the structural subtypes of CICs

To classify the cell-in-cell structures, we first divided the traditional CICs into five structural subtypes, including (a) partial, with more than 30% of the internalizing cells were enclosed, but not fully, by the outer cells; (b) one-in-one, with only one cell fully internalized, (c) two-in-one, with two cells were fully internalized; (d) in turn, a nested CICs with multiple cells sequentially internalized into neighboring cells; (e) complicated, a complex CICs generated by four or more cells (Fig. 3a). Considering the potential complexity, two kinds of breast cell lines including MCF7 and MCF10A were investigated, in which the total rate of CICs and its subtypes showed great discrepancy according to the manually labeling (Fig. 3b). In total, 17 pieces of MCF7 images and 85 pieces of MCF10A images were enrolled in this study, the cell number of each image ranged from 100 to 600, and from 30 to 200, respectively (Fig. 3c). The overall CICs rate of each image counted from 1% to 85% (Fig. 3d).

**Fig. 3.**
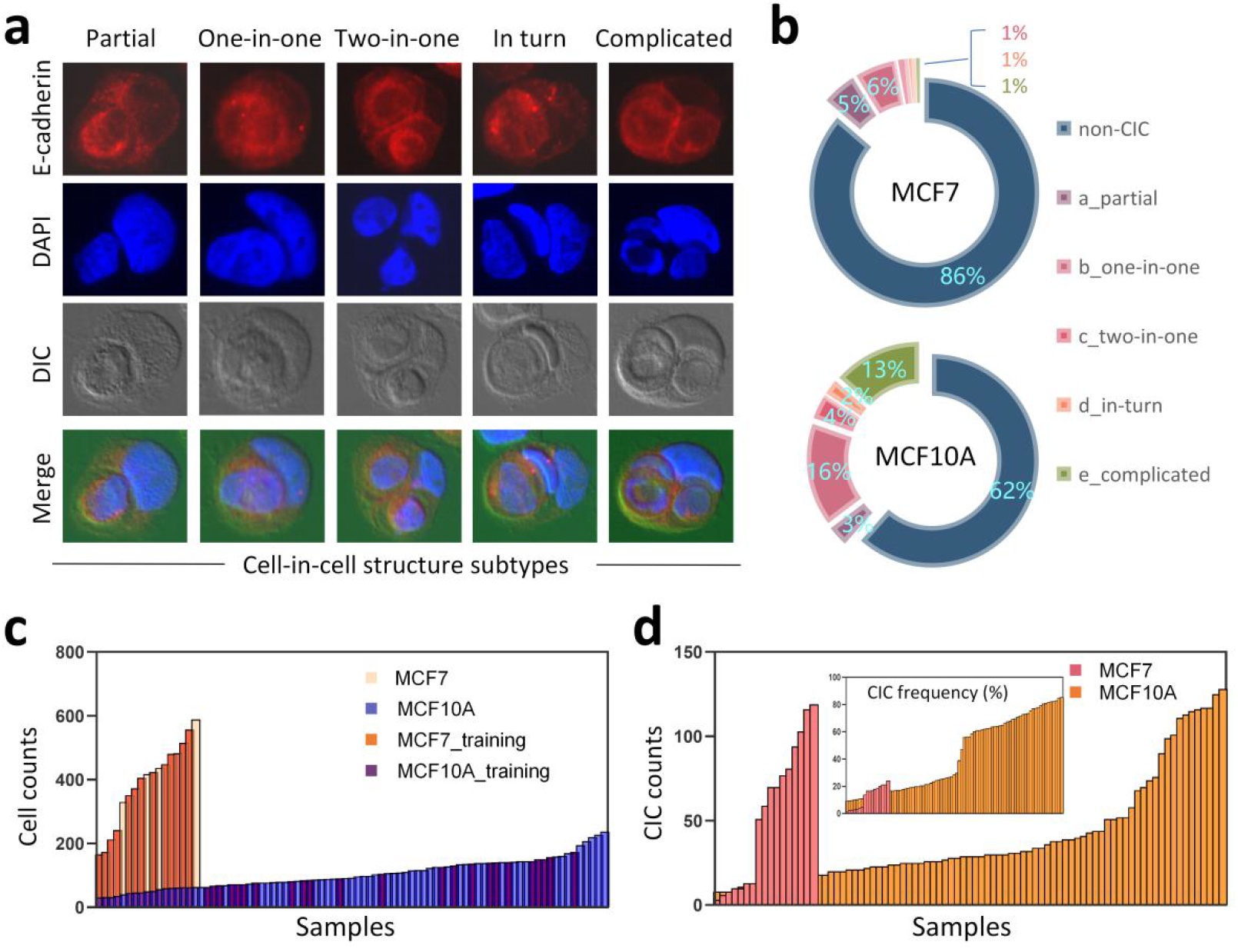
Image processing and cell candidate regions. **a** Representative images of five CICs subtypes. Cell membrane: E-cadherin in red, nucleus: DAPI in blue, and background is green. Original images are listed in supplementary Fig. S1a. **b** Percentage of different CICs subtypes for MCF7 and MCF10A cells. **c** The number of cell candidates extracted from each MCF7 and MCF10A image. Samples contained 17 pieces of MCF7 images and 85 pieces of MCF10A images. The columns in orange and purple represent for samples used in the training set. **d** The count and frequency of CICs for individual MCF7 or MCF10A image.

### Multi-Subtype classification achieved by the AIM-CICs

The obtained cell candidates were used to train ResNet101 model for the purpose of CICs recognition (Fig. S2a). Practically, we used 13 pieces of MCF7 images and 32 pieces of MCF10A images as the training set, which had 4026 MCF7 cells with a CICs rate of 11% and 3912 MCF10A cells with a CICs rate of 32% (Fig. S2b). Based on the morphological features of cell candidates, five subtypes of CICs were manually labeled for each cell candidate in the training and validation set. The distribution of each subtype of CICs showed remarkable discrepancy, as well as in the test set (Fig. 4a-b). To improve the practicality of the model, we defined a F-category from the non-CIC candidates. The F-category contains ambiguous structures that were hard to tell their identities by both experienced experts and AI algorithm, therefore, were generally removed from the sample counting (Fig S2c-d).

**Fig. 4.**
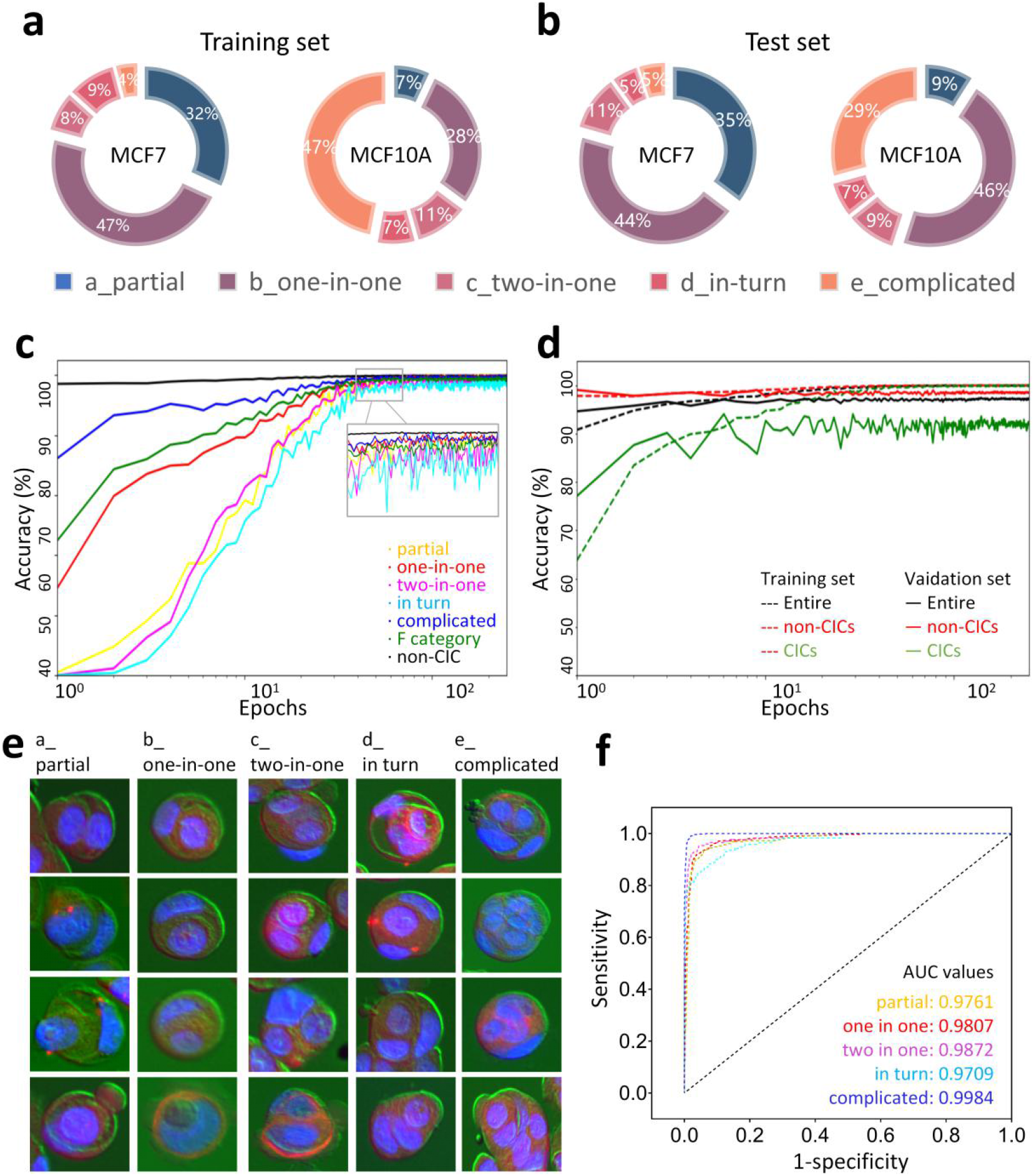
Training and testing process of the multiple classification. **a,b** The percentages of CICs subtypes in the training set (**a**) and test set (**b**). The CICs counts of MCF7 were 437 (**a**) and 340 (**b**), respectively. In the MCF10A samples, the CIC counts were 1269 (**a**) and 1948 (**b**), respectively. Supplementary figures associated: Supplementary Fig. S2. **c** The prediction accuracy of AIM-CICS for each subtype in a 250 epochs of learning process. **d** The integrated accuracy of AIM-CICS for CICs and non-CIC type in the training and validation set in a 250 epochs of learning process. CICs included the partial, one-in-one, two-in-one, in turn, and complicated type. Non-CIC referred to F-category and non-CIC. **e** Representative images of each CICs subtype predicted in the training set. **f** The ROC curves for each CICs subtype in the test set.

As shown in Fig. 4c, data training progressively increased the prediction accuracy to a considerable level for each subtype. In both training and validation sets, the comprehensive accuracy of integrated CICs (involving a, b, c, d, e types) and non-CIC type (including F category) revealed approving performance (Fig. 4d). Moreover, the AIM-CICs also exhibited impressive performance as indicated by the AUC of more than 0.97 for each CICs subtype (partial 0.9761, one-in-one 0.9807, two-in-one 0.9872, in turn 0.9709, complicated 0.9984) (Fig. 4e-f) in the test set. Additionally, for the low-quality images in the test set that displayed unclear cell regions and were eventually removed for further analysis, their recognition also reached an ideal AUC of 0.99 (Fig. S2e). Together, the AIM-CICs performed accurate recognition of CICs on independent datasets of MCF10A and MCF7 cells, suggesting the generalizability of this model.

### Visualization of morphological features and output

To better understand what the model learnt from the annotated data, we extract features from the output of network’s global average pooling layer and applied t-SNE to reduce dimension to 2D for visualization. For the training set, each group of cell samples represented independent clusters, except for cell candidates in the circled region (Fig. 5a). Backtracking the training data identified that these were candidates categorized into two subtypes due to erroneous manual annotation. Thus, the t-SNE-based clustering would be a visualized way for error-correction in recognizing CICs. For the test set (Fig. 5b), subtypes of CICs were clustered into close, but clearly distinct, regions, whereas F-category was neighboring to the area of non-CIC as expected. Moreover, following the comprehensive recognition under a specified confidence threshold, we were able to accurately locate each structure with a predicted value on the original images (Fig. 5c).

**Fig. 5.**
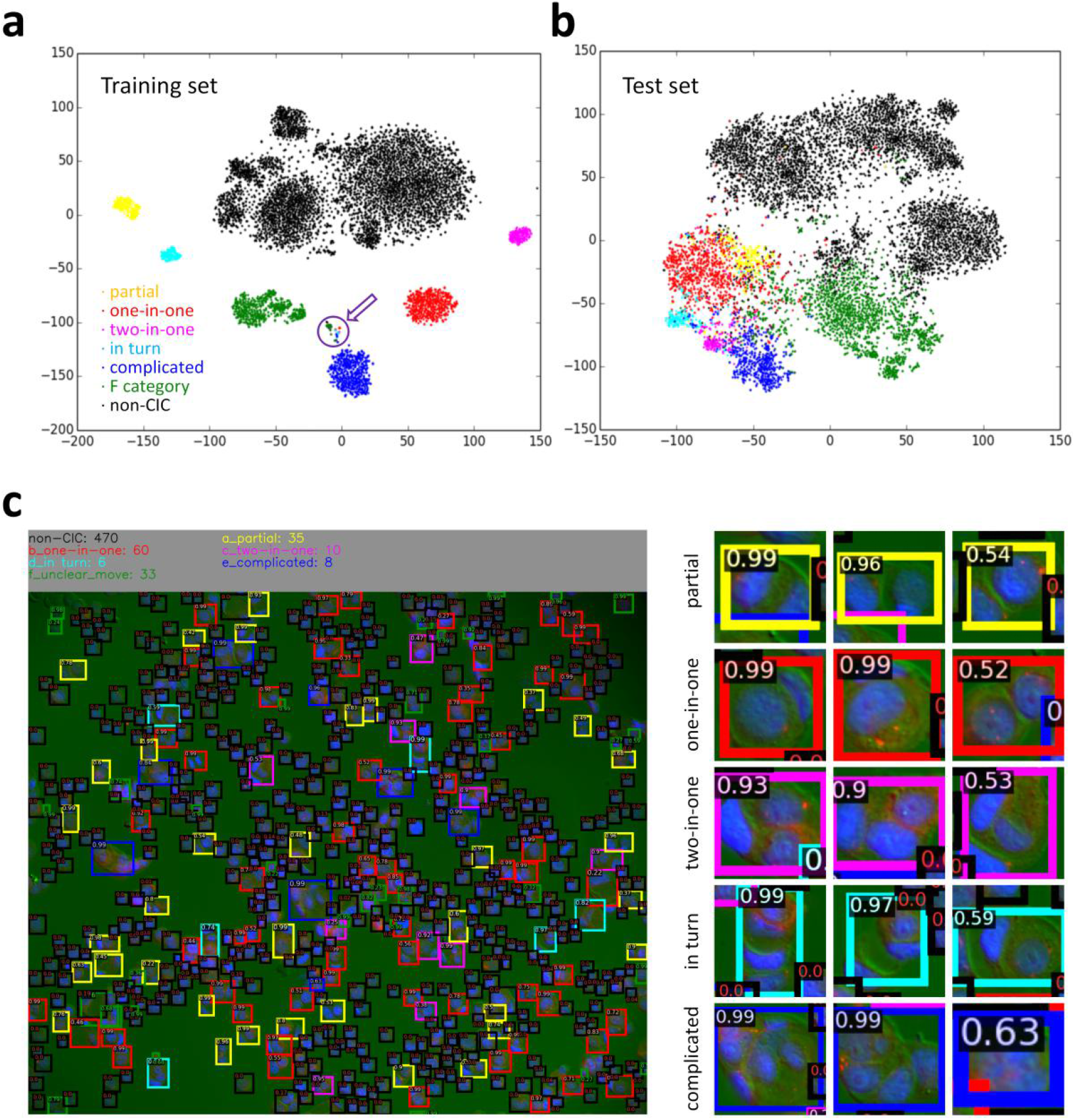
Visualization of sample features and output. **a,b** The two-dimension visualization of CICs subtypes in the training set (**a**) and the test set (**b**). The samples circled out were those predicted by the AIM-CICs to be miscategorized manually. **c** A representative image showing the recognition result of AIM-CICs. Colored Frames indicate structures in different categories, the predicated confidence scores were marked on the up-left corner of each structure. The structures in the right were cropped from the image in the left.

### Application of AIM-CICs in an experimental setup

To explore the potential implication of AI-based recognition of CICs in a biological context, we included a functional experiment as an example of subtype profiling. In this analysis, the confidence threshold was set to 0.2 for more informative identification (Fig. 6a). As the results showed, though all of the three truncations of ARHGAP36, a molecule identified to be a regulator of CICs formation in a screening study [20], resulted in impaired formation of CICs, the alterations of CICs subtypes were rather different (Fig. 6b-c). While the truncated GAP36 (1-194) had little impact on the formation of partial CICs (Fig. 6b-d), the majority of CICs were in completed form (including all CICs subtypes except for the partial) in cells expressing the truncated GAP (118-194) or GAP (195-395) (Fig. 6b-c), suggesting that the N-terminal region (1-117) of ARHGAP36 might function to slow down the process of cell internalization. Meanwhile, the C-terminal region of ARHGAP36 was likely to be responsible for the closing step of CICs formation as evidenced by comparable formation of completed CICs between control and GAP (195-395)-expressing cells (Fig. 6b and 6e-h). Moreover, the GAP (118-194) seemed to be the major region that drives cell internalization as it promoted the formation of completed CICs at a rate comparable to the GAP (1-194) region. Furthermore, though the N-terminal region might negatively regulate the speed of CICs formation, it did function positively to promote cell internalization as its truncation significantly reduced the formation of both partial and total CICs (Fig. 6b-d). Thus, the AIM-CICs algorithm allows us, for the first time, to accurately dissect the impacts of different domains or molecules on CICs formation in a heretofore underappreciated resolution.

**Fig. 6.**
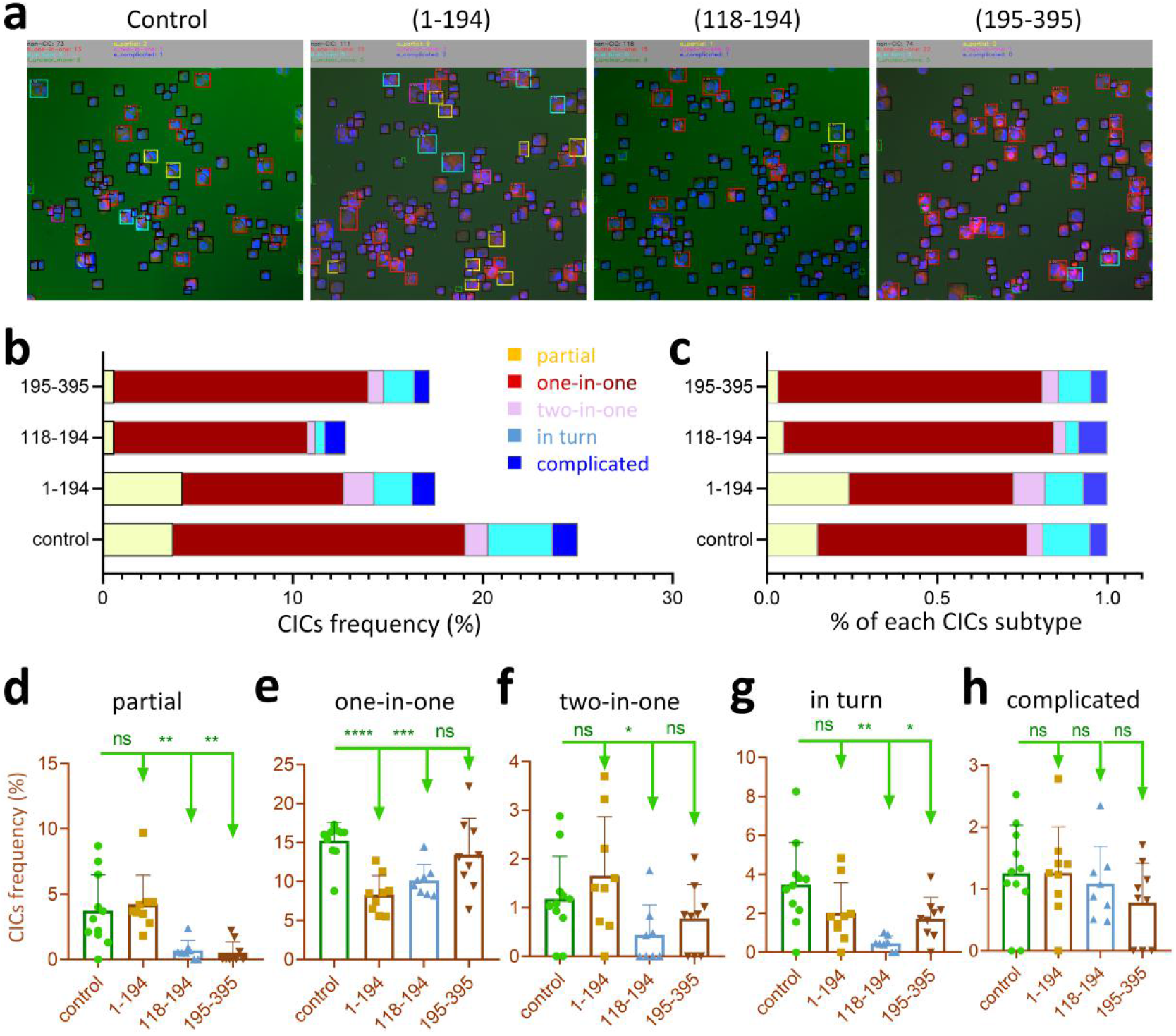
Analysis of CICs subtypes in an experimental setup by AIM-CICs. **a** The visualized recognition results of AIM-CICs in an experiment, where MCF10A cells expressed the empty vector (control) and three truncated mutants of ARHGAP36 (1-194, 118-194, 195-395), respectively. **b,c** Graphs show the absolute frequency (**b**) or relative frequency (**c**) of subtyped CICs in MCF10A cells expressing different ARHGAP36 mutants. (n = 934 cells for control, 1060 cells for 1-194, 1392 cells for 118-194, 852 cells for 195-395). **d-h** The frequencies of subtyped CICs in MCF10A cells expressing different ARHGAP36 mutants. Data were shown as box-plots with means and individual data points. * for *p* < 0.05, ** for *p* < 0.01, *** for *p* < 0.001 (two-tailed Student’s t test).

## Discussion

Fluorescent microscope images recorded the cellular structures such as CICs, but inevitably provided a great number of morphological variations. To provide recognition with sufficient accuracy and potentially featured insights, we, for the first time, explored the application of Convolution Neural Network (CNN) in the profiling of subtyped CICs formed during entosis, a non-apoptotic cell death process occurred via cell-in-cell invasion [11]. Based on the tons of images accumulated from previous studies, we developed the AI-based identification algorithm AIM-CICs, which was trained with distinct illumination, textures, and density, in order to deliver an optimal performance in cell region detection and multiple subtypes classification, despite of the unseen perturbations.

In the proposed system, we set up two tasks, of which, a classic 1-class object detection model was formulated to find cell regions as the first task, followed by multi-class object recognition as the second task. Comparing with the traditional end-to-end manner, i.e., to train a multi-class detection model with different kinds of cells marked simultaneously, our model of separated detection will achieve the flexibility for the raw samples to be recategorized and repurposed. In AIM-CICs developed in this study, the second task included a well-trained 7-category classifier (5 CICs subtypes plus one non-CICs and one F-category) to define the multiple subtypes of CIC structures, which is compatible with the cell candidates from the first step. This two-step algorithm is also advantageous in debugging the possible mechanisms leading to inferior final prediction outcomes, as each step could be optimized separately. Meanwhile, this two-step algorithm may fall short of efficiency (speed) as compared to the end-to-end multi-class detection model which could utilize a shared feature extraction backbone.

Among all the defined cell death programs, CICs-mediated death is unique in that it can only be accomplished with the involvement of at least two cells, but not one cell in other programs like apoptosis, necrosis and the forth [30]. Therefore, mechanistic study is a challenging task for the field of CICs-mediated death, which was further complicated by the fact of lacking a reliable biochemical marker. Current studies on CICs relayed on the morphology-based binary quantification, that is, CICs or non-CICs. Here, CICs were usually defined as structures with more than 1/2, or 2/3 in some studies, of the inner cell body being internalized/enclosed by the outer cell. This oversimplified quantification of CICs, did move the field forward over the past decade, however, provided rather coarse information over a more complicated process [22, 31]. CICs formation is a stepwise process that could be empirically subdivided into three major stages: 1) the early initiation stage from cell-cell contact to about 1/3 of the inner cell body being internalized, this stage was primarily driven by cell-cell adhesion and assisted by cytoskeleton remodeling; 2) the middle internalization stage covering the whole process of cell internalization that was primarily driven by active actomyosin contraction within the inner cells, and coordinately assisted by the outer cells; 3) the final closing stage that may involve in tail cutting and membrane fusion, this a process rarely being investigated largely because it is technically challenging. Furthermore, CICs formation is a dynamic process that may have multiple cells, either sequentially or simultaneously, form a complicated structure that may contain more than one cell inside (Fig. 3a). The regulation of this feature is completely unknown for the field yet, but might be conceptually feasible as it was reported in phagocytosis that the number of corpses engulfed by a phagocyte was genetically controlled [32]. Taking these two factors (stage and cell number) into account will produce an even higher dimensional complexity, which, however, was missed from the traditional analysis by the binary quantification. The implantation of AIM-CICs enables us to make a more sophisticated description of CICs phenotypes, which would help identify finer molecular control. For example, though expression of the GAP (195-395) domain did not influence the frequency of simple CICs, where only one cell was enclosed (one-in-one in Fig. 6e), it did result in significantly reduced formation of complex CICs, where more cells were enclosed by one outer cell (two-in-one and in-turn in Fig. 6f-g). These results suggested that the truncated N-terminal domain may facilitate the internalization of multiple cells to form complex CICs, which warrants further functional validation.

In addition to mechanistic investigation, AIM-CICs is also promising in enabling high-content based screening for therapeutic compounds that target CICs formation considering their pivotal roles in multiple biomedical processes such as cancer [1]. While high throughput screening generally relies critically on a reliable biochemical marker that is currently unavailable to CICs formation, the related systemic screening, which would be labor-intensive and time-consuming if worked out by manual annotation, had yet been reported. Empowered with AIM-CICs and high-content microscopy, the systemic screening would be feasible in the near future.

## Acknowledgements

We thank Dr Lulin Zhou, Xiaoyi Jiang, He Ren, Yichao Zhu, Yuqi Wang, Lihua Gao, Zhaolie Chen, and the members of the Sun lab for the constructive discussions and assisting manual labeling.

## Author contributions

Concept and design: QS; CNN Model Training and Analysis: WZ; Data collection: MT, ZBN, CXW, and BZR; Data interpretation: QS, WZ and MT; Figures: MT, YS, WZ and QS; Manuscript: MT, QS and WZ, with input from ZBN, CXW, BZR, YZ, BZ, QQL, YZ, HYH, FXZ, and HPS. Funding acquisition: QS, HYH, HPS, and MT. All authors have read and approved the final manuscript.

## Ethics statement

This study did not involve ethical approval.

## Materials and methods

### Image processing and softwares

An entire dataset involving 17 pieces of MCF7 images and 85 pieces of MCF10A images were obtained from Sun’s lab. As detailed protocol described [33], the fluorescently labeled cells were necessary to be stained with discrepant colors for each cell components, such as, red for cytomembrane (E-Cadherin, 1:200, BD Biosciences, 610181) with secondary antibodies Alexa Fluor 568 anti-rabbit (1:500; Invitrogen; A11036), and blue for cytoblast (DAPI, Sigma D8417). Random fields were taken under corresponding channels of laser lights through fluorescent microscopy (Nikon Ti-E microscope, Nikon NIS-Elements AR 4.5 software), along with bright color for the background. For algorithm performing, each sample with three single-channel images was transformed into an RGB format with value rescaled to 0 - 255. Softwares used and algorithms developed in this study include: Python (http://www.python.org./); PyTorch (https://pytorch.org/); VIA Annotation Tools (https://www.robots.ox.ac.uk/~vgg/software/via/); Detectron2 (https://github.com/facebookresearch/detectron2).

### Cell region labeling and candidates extraction

After acquiring the processed images, we manually annotated the cell regions through VGG Image Annotator (https://www.robots.ox.ac.uk/~vgg/software/via/). Based on the annotated images, a classic 1-class object detection task was carried out for cellular morphological learning. The model we used is a Faster-RCNN [28] network with ResNet-50 [29] backbone. Since the original resolution of microscopic image is 2160 × 2560 which is too large for Faster-RCNN training, we first split each image into 4×4 grids, then follow the common practice to train the model. For data augmentation, we use random flip, random rotation, and random scale to expand diversity of data. As for other hyper-parameters, we set batch-size to 24 and iterate 50000 steps using SGD optimizer with momentum 0.9. As the output of the Faster-RCNN network, the patches of detected cell regions were exported as candidate sequences for further steps.

### Manual classifications of cell-in-cell structures

The manual definition of cell-in-cell structural classification primarily included bipartite-class, CICs and non-CICs. CICs category were further subdivided into 5 subtypes, including (a) partial, with more than 30% of the internalizing cells were enclosed, but not fully, by the outer cells; (b) one-in-one, with only one cell fully internalized, (c) two-in-one, with two cells were fully internalized; (d) in turn, a nested CICs with multiple cells sequentially internalized into neighboring cells; (e) complicated, a complex CICs generated by four or more cells. To refine the output results, we added a F-category among non-CICs, which was defined as unclear or not sure for the cell recognition and needs to be removed for the quantitative analysis. The cell candidates involved in the training set were verified together by an expert group consisted of 6 members in the lab.

### Multiple classification model

We used the ResNet101 model as our classifier and the input size was set to 224. Since this model could take the detection model’s output as input, we cropped cell samples using detection model and manually labeled them with corresponding cell types. During training, each sample was first padded to square and then resize to 224 × 224. Both horizontal and vertical random flip were performed. We trained our model for 250 epochs with batch size of 32, using SGD optimizer with learning rate of 0.001 and momentum of 0.9. To alleviate overfitting, a dropout layer with *p* = 0.25 was set right before feature went into the final fully connected layers. To choose hyper-params, we kept 20% samples as validation set. Eventually, the prediction results could be visualized on the original image with detected cell region and a predictive score of CICs, as well as in the output folders of each cell type.

Importantly, when applying our model for inference, the test samples should be pad and resize in the same way as training. Our model is a 7 classes classifier, and it outputs a 7 elements vector representing the probability for the test sample to belong to each type. Traditionally, the predicted type should be the type with maximum probabilities. In practice, to increase precision, we predict cells that have predicted probability lower than 0.2 as non-CIC, even if the non-CIC probability is not the maximum for it. For example, if the predicted output is [0.1,0.18,0.12,0.15,0.15,0.13,0.17] (for a, b, c, d, e, f, non-CIC), we will use non-CIC as model’s prediction. Ultimately, we could output the classifications into specific folders of each cell type, and obtain the visualized results that marked with individual colors on the original image.

### Performance analysis of detection model

In deep learning community, the most common metric used for quantitatively comparing detection models’ performance is mean average precision (mAP), as proposed in [34] and [35]. However, since our work mainly focused on multi-type CICs classification instead of general object detection technique, we reported our detection result in a more practical recall/precision manner. In detail, we kept detection model’s output instances with confidence > 0.1 as model’s prediction and calculate metrics at two different Intersection over Union (IoU) thresholds 0.5 / 0.1. Under IoU threshold 0.5, the model must output an accurate prediction box to get a match, while 0.1 requires only loosely overlapping.

### Features visualization

To better understand what the classification model learns from labeled samples, we extracted features from each cell sample and visualize them in a 2D space. The feature we used is the output of network’s global average pooling layer, which is right before the final classification layer. This 2048D feature is the deepest and the most semantic so it can represent the information extracted by the network from a corresponding input image. To visualize these 2048D features, we uses the t-SNE method for dimensionality reduction to transform each feature to 2D [36]. t-SNE is a popular method for visualizing high-dimension data since it can keep most of the original data structure during dimensionality reduction.

### Evaluation criteria for classification models

The output of classification model was evaluated by the universal criteria, such as, sensitivity (Se or recall), specificity (Sp), precision, the receiver operating characteristic (ROC) curve, and the area under ROC curve (AUC). The equations were listed as follows:

1. Se (recall) = TP / (TP+FN)
2. Sp = TN / (TN+FP)
3. Precision = TP / (TP+FP)

True positive (TP) stands for the accounts of positive CICs which are correctly recognized as positive CICs. False positive (FP) stands for the number of negative CICs that are incorrectly recognized as positive CICs. False negative (FN) stands for the accounts of positive CICs which are incorrectly recognized as negative CICs. True negative (TN) stands for the number of negative CICs correctly recognized as negative CICs.

### Statistical analysis

Categorical data are expressed as frequencies (%) and were tested by a two-tailed Student’s t-test. P values were calculated by Excel or GraphPad Prism software. The level of significance was set at *p* < 0.05.

---

**Fig. S1.**
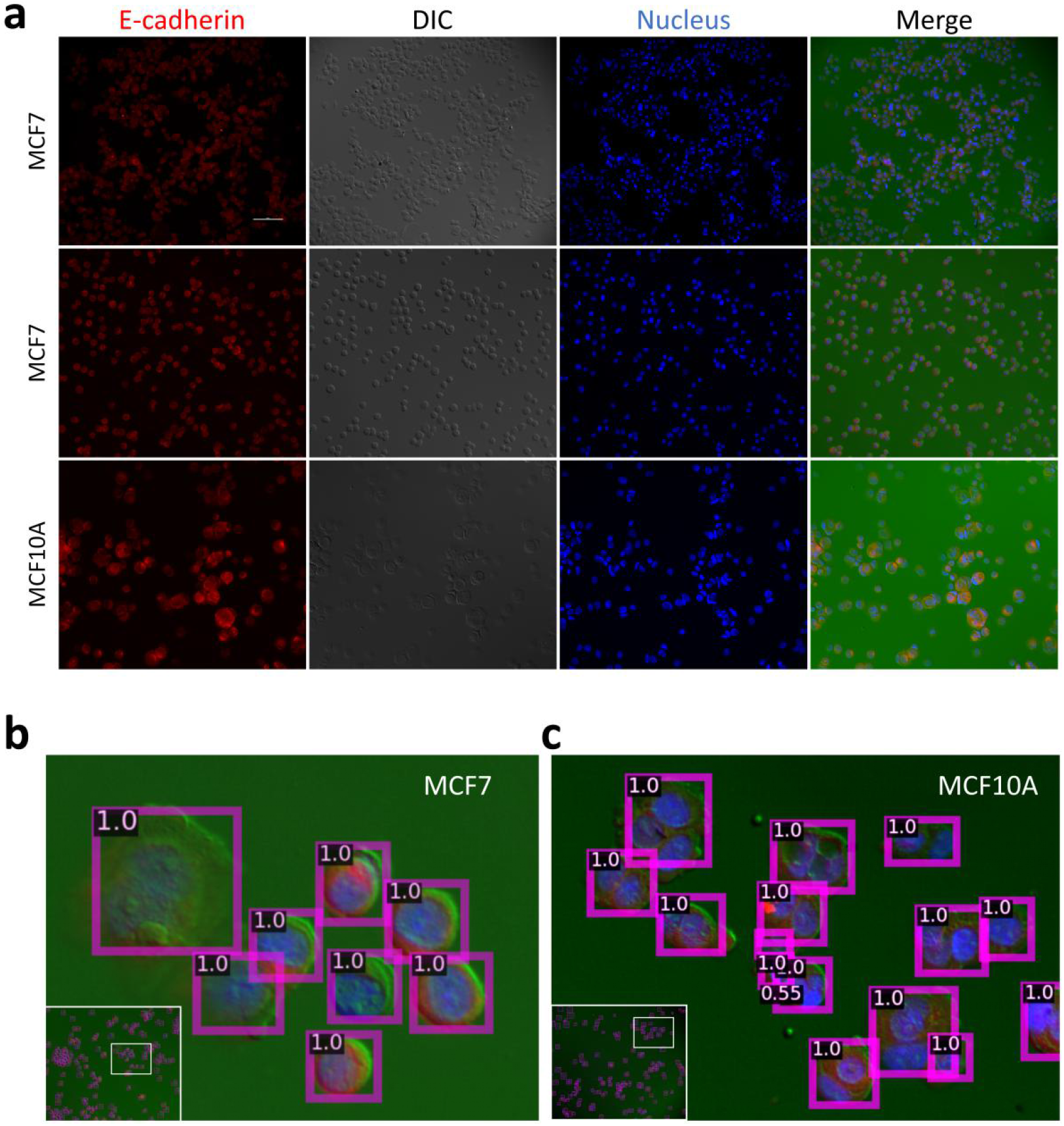
Format transformation and cell region detection. **a** Representative images of different brightness in RGB format for MCF7 and MCF10A cells, respectively. Cell membrane: E-cadherin in red, nucleus: DAPI in blue, and background is green. The intensity value was rescaled to 0 - 255. **b,c** Representative images of MCF7 (**b**) and MCF10A (**c**) samples predicted by the cell region detection model. Each cell region was indicated by a box with a predicted confidence score.

**Fig. S2.**
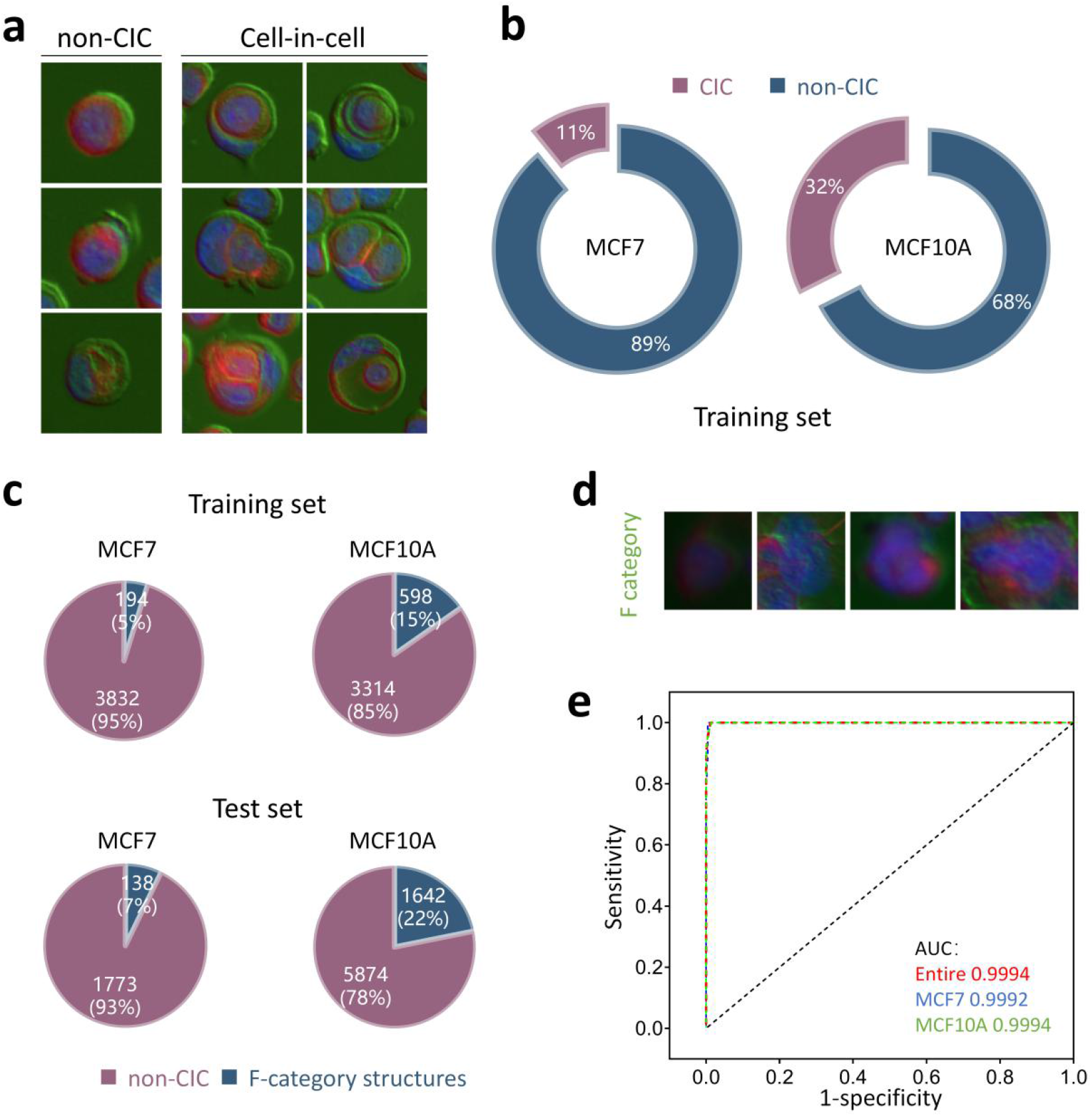
Binary subtyping by the AIM-CICs. **a** Representative images for CICs and non-CICs. **b** The quantification of CICs for MCF7 and MCF10A cells in the training set. **c** Percentage analysis of F-category relative to the non-CIC category in training set and test set. **d** Representative images of F-category, images belonging to this category were unclear and hard to be classified into other categories. **e** The ROC curves for the F-category in the entire dataset, MCF7 group, and MCF10A group of test set, respectively.

